# ESPWA: a deep learning tool to inform precision-based use of endocrine therapy in resource-limited settings

**DOI:** 10.1101/2025.08.27.672012

**Authors:** Dagoberto Pulido-Arias, Rebecca Henderson, Milit Patel, Christophe Millien, Joarly Lomil, Marie Djenane Jose, Gabriel Flambert, Jean Bontemps, Emmanuel Georges, Alekhya Gunturi, Palak Shah, Tiago Goncalves, Jayashree Kalpathy-Cramer, Elizabeth Gerstner, Seth A. Wander, S. Joe Sirintrapun, Dennis Sgroi, Jose Jeronimo, Philip E. Castle, Kenneth Landgraf, Ali Brown, Temidayo Fadelu, Lawrence N. Shulman, John Guttag, Dan Milner, Jane Brock, Christopher P. Bridge, Albert E. Kim

## Abstract

Immunohistochemistry for estrogen receptor (ER) expression is often unavailable in low-and-middle-income countries (LMICs), leading to empiric use of endocrine therapy (ET) and unnecessary toxicity in ER-negative patients. To address this unmet need, we developed **ESPWA,** a deep-learning model trained on 3448 H&E slides and tissue-matched ER status from breast cancer patients treated at Zanmi Lasante (ZL), Haiti. A model trained on The Cancer Genome Atlas (TCGA) exhibited substantial domain shift when applied to the ZL cohort, with AUROCs dropping from 0.846 on TCGA cross-validation to 0.671 on the ZL cohort. In contrast, ESPWA demonstrated improved performance on ZL cross-validation (AUROC=0.790; p=0.005). In an independent test set of 134 Haitian patients with parallel slides prepared and scanned in Mirebalais Hospital (Haiti) and Brigham and Women’s Hospital, ESPWA was robust to variations in slide preparation, quality, and scanners, achieving AUROCs of 0.794 on BWH-prepared WSIs and 0.805 on Mirebalais-prepared WSIs. Prospective studies using ESPWA are underway in sub-Saharan Africa to evaluate its utility in informing precision-based use of ET.

## INTRODUCTION

Breast cancer is the most common cancer type and leading cause of cancer mortality for females worldwide^1^. This mortality disproportionately affects patients in LMICs, rather than those in high-income countries (HICs). Despite recent advances in hormonal and targeted therapy that render early-stage breast cancer increasingly curable^2^, patients in LMICs continue to experience delayed diagnoses and suboptimal tumor-directed therapy^3^. In Haiti, one of the most resource-limited settings in the world, the challenge is especially dire. As only five pathologists serve the entire country of nearly 12 million people, timely and complete pathologic assessment of tumor tissue is nearly impossible. A delayed diagnosis of cancer leads to a higher likelihood of the patient presenting in the metastatic stage, which in turn decreases life expectancy. The mortality-to-incidence ratio for Haitian breast cancer patients is over 60%^4^, compared to just 2.3% in the United States^5^. This stark disparity underscores an urgent need for innovative methods that enable delivery of high-quality cancer care in low-resource settings.

In clinical workflows, pathologists perform a manual analysis of H&E-stained slides, which are widely available in HICs and LMICs. However, as a typical H&E slide contains hundreds of thousands of cells, it is impractical for a human to ingest, analyze, and report this vast amount of data expeditiously. To this end, DL applications in digital pathology have emerged as powerful tools to enhance the efficiency and yield of pathology workflows. Furthermore, recent studies have demonstrated DL’s potential to identify subtle spatial patterns of cell populations^6,7^ and individualized therapeutic insights, thus highlighting its potential for precision oncology.

Given efficacy observed with endocrine therapy for ER-positive breast cancer^8^, a timely assessment of ER status is arguably the most critical element of the diagnostic-therapeutic pathway for breast cancer. However, immunohistochemistry (IHC), the gold standard for measuring ER expression, is often unavailable in LMICs and dependent on donated services from academic medical centers and medical equipment companies in HICs, thus resulting in delays on the order of months. Therefore, breast cancer patients in Haiti and other LMICs are often empirically treated with ET^9^, which subjects ER-negative patients to ineffective therapy and unnecessary toxicity^10^.

To address this clinical need, we developed **ESPWA** (**E**strogen Receptor **S**tatus **P**rediction for Haitian patients using deep learning-enabled H&E **W**hole Slide Imaging **A**nalysis), a DL model that infers quantitative ER expression from H&E-stained WSIs. Although ER status is not routinely assessed from H&E slides in clinical practice, prior DL models have demonstrated the feasibility of inferring ER expression from H&E WSIs^11–13^. However, these models have been trained on publicly available datasets of H&E WSIs primarily sourced from HICs. Given intrinsic differences between HIC and LMIC datasets, such models may not generalize well to the LMIC population. There are notable differences in tissue quality and fixation protocols between HICs and LMICs^9,14^. In addition, publicly available datasets such as The Cancer Genome Atlas (TCGA) are largely derived from patients of Caucasian descent^15^, which differs from the predominantly Black population in Haiti and other LMICs. These differences are relevant for model generalizability, given a higher incidence of aggressive histological features and poorly differentiated tumors in the Black population compared to Caucasians^16–18^.

In this proof-of-concept study, we demonstrate the utility of DL to bridge the diagnostic gap in resource-limited settings by extracting individualized therapeutic insights from routine H&E WSIs. To train ESPWA (**E**strogen Receptor **S**tatus **P**rediction for Haitian patients using deep learning-enabled histopathology **W**hole Slide Imaging **A**nalysis; “Hope” in Haitian Creole), we curated a dataset of H&E WSIs and tissue-matched ER status from breast cancer patients treated at Zanmi Lasante (ZL), the largest provider of cancer care in Haiti. All tissue samples within our training set were fixed and embedded locally, ensuring that our dataset reflects the variability in tissue quality observed in LMICs. Next, using an independent test set from 134 Haitian breast cancer patients with H&E slides that were independently cut, stained, and scanned at both Mirebalais Hospital (Haiti) and BWH, we demonstrate that ESPWA achieves robust performance in assessing ER status across variations in slide quality and scanners and across tissue types and histological subtypes. To build upon this work, we have initiated prospective clinical trials in ZL and partner sites in eight African countries to evaluate the utility of ESPWA in informing precision-based use of ET in resource-limited settings. More broadly, this work serves as a scalable framework for employing DL to interrogate tumor biology across large heterogenous populations.

## METHODS

### Dataset curation

We curated three distinct cohorts of H&E whole slide images (WSIs) and tissue-matched ER status to develop and validate our models. To establish a baseline for model trained on high-income country (HIC) data, we used The Cancer Genome Atlas – Breast Invasive Carcinoma (TCGA) dataset. This cohort comprised 1085 H&E WSIs from 1073 patients with primary breast tumors. Each WSI had tissue-matched ER status derived from IHC.

ESPWA’s training set consisted of 3448 H&E WSIs from 1808 breast cancer patients from Hôpital Universitaire de Mirebalais, the teaching hospital of Zanmi Lasante (ZL) in Mirebalais, Haiti. This cohort included 2933 primary tumors and 515 lymph node metastases. Tissue blocks were created in ZL. Sectioning and staining with H&E were performed at the Brigham & Women’s Hospital (BWH). ER status for all ZL cases was determined with IHC at BWH. This IHC-derived ER status served as the ground truth label for the ZL dataset. All H&Es were digitized using the KFBIO KF-PRO-005 scanner at 40x magnification.

To evaluate model generalizability across laboratory protocols and scanning hardware, we curated an independent test set of 134 Haitian breast cancer patients with IHC-confirmed ER status. This cohort employed a paired design: for each patient, parallel sets of H&E slides were independently cut, stained, and scanned at both Mirebalais Hospital (MoticEasyScan) and BWH (KFBIO KF-PRO-005), totaling 290 WSIs. This strategy allowed for a direct head-to-head comparison of model performance on slides processed in a resource-limited settings versus those processed at a HIC academic center.

We evaluated image artifacts in the Mirebalais and BWH cohorts using the GrandQC computational pipeline. GrandQC performs automated image segmentation to generate spatial maps (GeoJSON masks) that physically outline regions containing slide preparation and scanning artifacts. Our analysis focused on four specific artifact types identified by the pipeline: tissue folds, dark spots/foreign bodies, edge/bubble artifacts, and out-of-focus regions. From these spatial masks, we calculated the total area of each artifact type per slide. We then normalized these areas as a percentage of the total valid tissue area. Because these artifact percentages followed non-normal distributions, we applied a non-parametric two-sided Mann-Whitney U test to assess the statistical differences in median artifact proportions between the Mirebalais and BWH-prepared cohorts.

### WSI Processing

For both the TCGA and ZL datasets, quality control and tissue segmentation were performed as per prior literature^19,20^. In addition, we performed label-preserving data augmentation to extracted image patches (**Figure 1A**) on the training set. Further details are provided in the *Supplementary Methods*.

**Figure 1:**
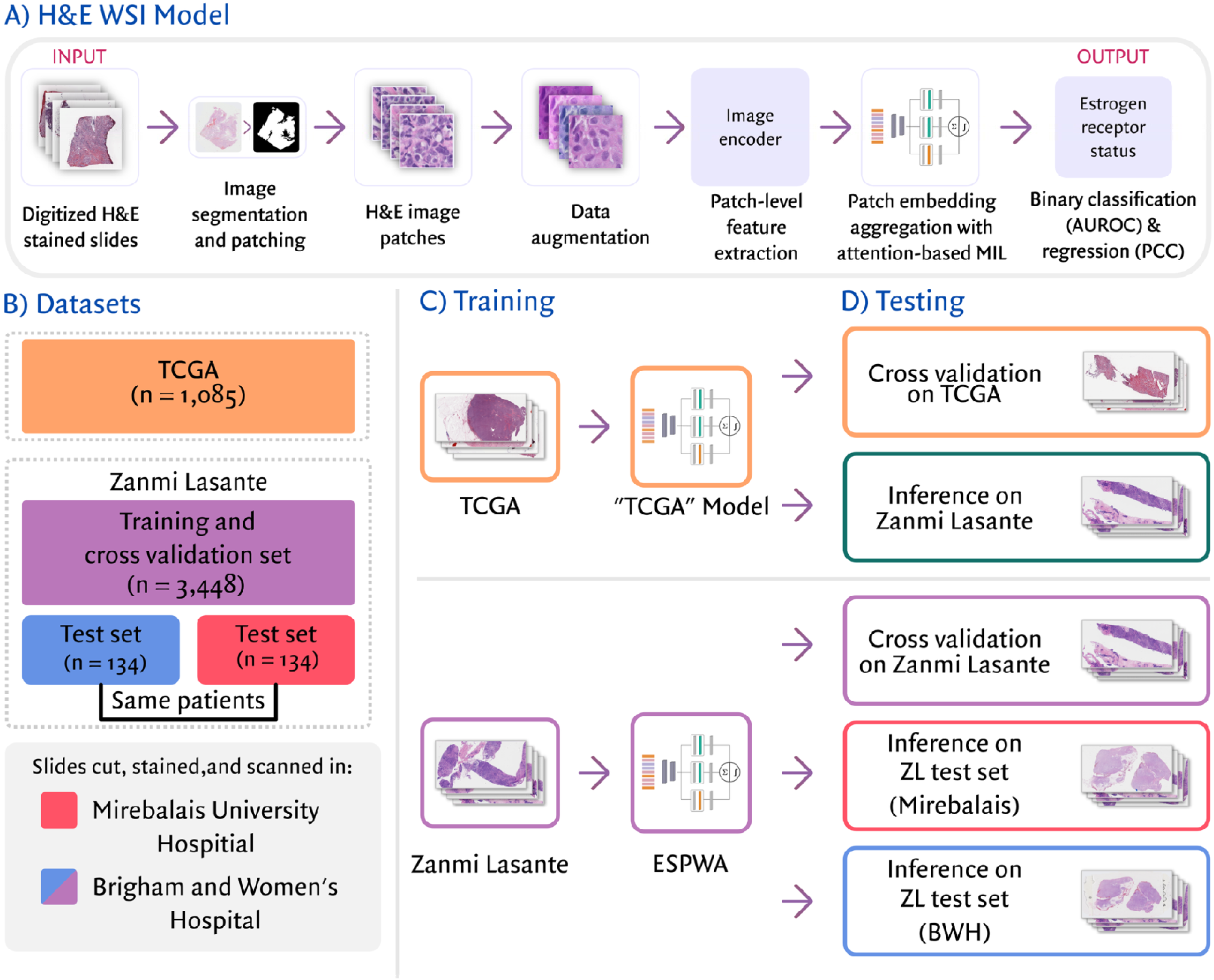
Overview of ESPWA development and evaluation. A) H&E WSI Model. Digitized H&E WSIs are segmented into patches and encoded via a frozen image encoder. An attention-based multiple-instance learning aggregator integrates these patch-level feature vectors for binary classification and regression of ER status. Ground truth for all H&E WSIs was defined by tissue-matched IHC-derived ER expression. B) Datasets. Models were developed using TCGA (n = 1085) and Zanmi Lasante (n = 3448) cohorts. A paired test set (n = 134 Haitian patients) was used to assess robustness to technical variability within LMIC settings, where parallel slides from the same patient tissue blocks were independently processed at Mirebalais Hospital (Haiti) and BWH (United States). C) Model training. Two distinct models were developed: a “TCGA” model and ESPWA. ESPWA was trained on the ZL dataset to capture biological and technical nuances specific to the Haitian population and pathology labs. D) Testing. Performance for the “TCGA” model and ESPWA was assessed through 10-fold cross validation. To evaluate for domain shift, the “TCGA” model was tested on the ZL cohort. ESPWA was first evaluated using the internal ZL test folds. Following this, ESPWA was validated on the paired BWH and Mirebalais-prepared test sets to demonstrate invariance to laboratory-specific technical variability and WSI quality.

### Model training

We trained two models to predict ER status from H&E WSIs: one trained on the TCGA dataset for training, and the other trained on the ZL cohort (**Figure 1**). These models were developed using 10-fold cross-validation, with data partitioned at the patient level (70% for training, 15% for validation, and 15% for testing). We assessed the impact of domain shift across the TCGA and ZL datasets by evaluating the TCGA model on the ZL dataset.

Both the TCGA model and ESPWA shared an architecture consisting of (1) a frozen, pre-trained image encoder using the publicly available weights from the CONtrastive learning from Captions for Histopathology (CONCH) model^20^ that converts high-resolution image patches into a 768-dimensional feature vector; (2) a trainable multiple instance learning (MIL) attention aggregator that integrates all encoded patches from a slide into a global slide-level feature representation following the clustering constrained attention multiple instance learning (CLAM) architecture^19^; (3) a trainable fully-connected classifier head with a sigmoid activation that outputs the probability of a patient’s ER status based on the slide-level embedding. The TCGA model and ESPWA quantified ER expression using binary classification: ER-negative or ER-positive. As the ZL cohort also had quantitative ER expression scores, we also trained a separate regression model to assess a quantitative measure of ER expression from the H&E WSI.

Further details about model architecture, hyperparameters, and hardware used for model training are included in *Supplementary Methods*.

### Analysis of model performance

For binary classification, we evaluated model performance with accuracy, sensitivity (recall), positive predictive value (precision), area under the receiver operating characteristic curve (AUROC), and the area under precision-recall curve (AUPRC). For regression, we evaluated model performance with the Pearson’s correlation coefficient (PCC) and intra-class correlation coefficient (ICC). All primary performance metrics were calculated using 10-fold cross validation. To estimate variability, 95% confidence intervals were computed via bootstrapping with 1000 re-samples across the test folds. Performance comparisons across different models and subgroups were compared using the DeLong test. On the independent test set (n = 134), ESPWA was validated by comparing the AUROC achieved on Mirebalais-prepared WSIs versus BWH-prepared WSIs, with 95% Cis and significance testing performed as described above.

We also conducted a decision curve analysis following a previously published framework^21–23^ to evaluate the benefit of the model to support clinical decision making under a range of different preferences. Since the current standard practice in Haiti is to treat all patients with ET, we use the “interventions avoided” variant that measures net benefit in terms of a balance between the net decrease in false positives and net increase in false negatives compared to the “treat all” strategy that considers all patients as ER-positive.

### Interpretation of model predictions via attention heatmaps

To interpret the relative importance of different regions in a WSI to the final WSI-level prediction of the model, we created heatmaps with colors corresponding to attention weights of each patch token used to calculate the slide-level embedding^19^.

## RESULTS

### Substantial domain shift between TCGA and ZL datasets

Consistent with previous studies^11,13^, our TCGA-trained model demonstrated strong performance in predicting ER status using H&E WSIs from a predominantly Caucasian population. Our TCGA model achieved an accuracy of 0.850 (95% CI: 0.827-0.879) and an AUROC of 0.846 (95% CI: 0.831-0.879) on the held-aside TCGA test sets (**Figure 2**). Next, to evaluate the ability of our TCGA model to generalize to Haitian patients, we ran inference using the ZL dataset. We observed a marked decrease in performance, with an accuracy of 0.600 (95% CI: 0.590-0.611) and an AUROC of 0.671 (95% CI: 0.662-0.675, **Figure 2**).

**Figure 2:**
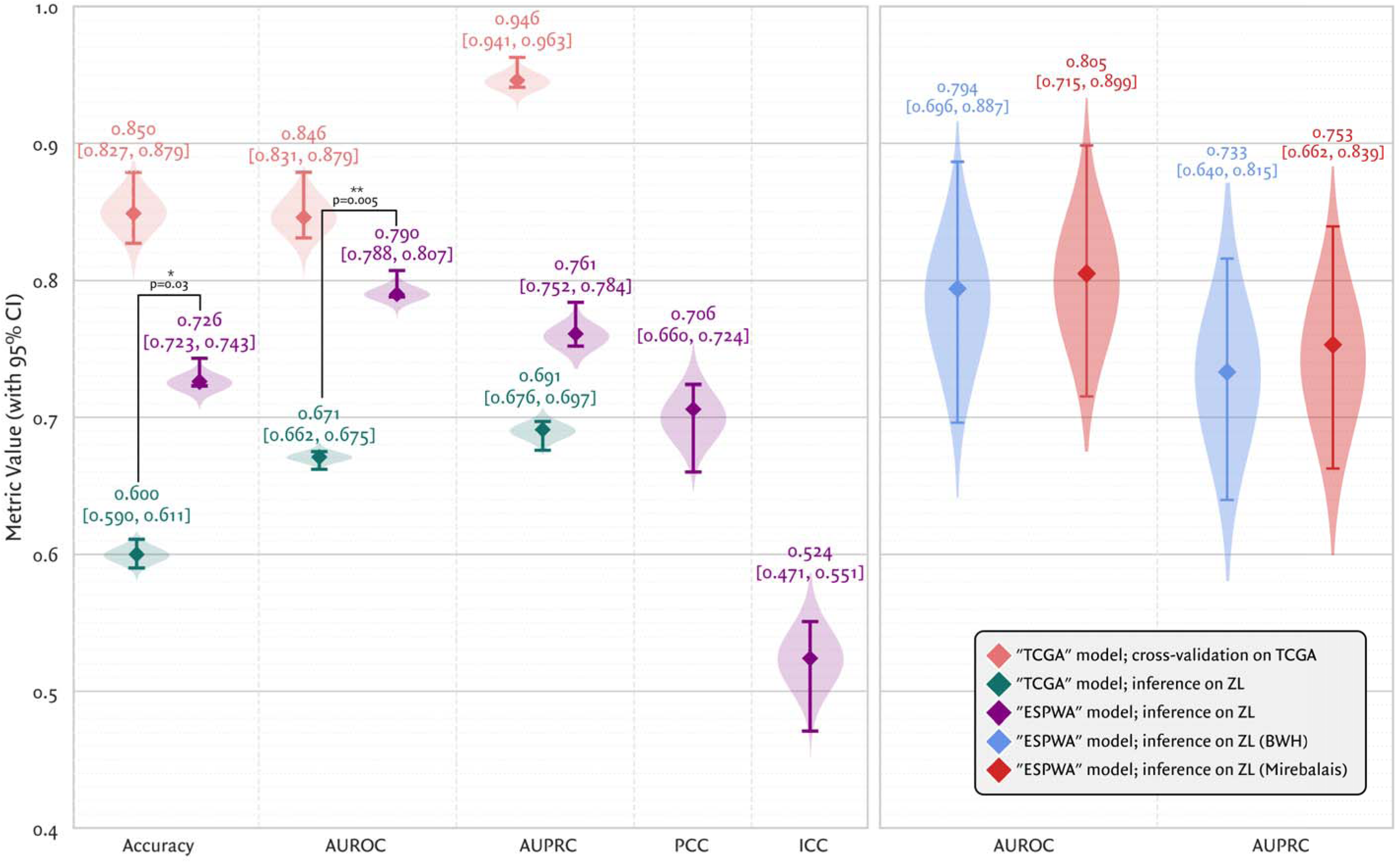
“TCGA” model and ESPWA performance across cross-validation and test sets. Violin plots display distributions of performance metrics (diamond = median; whiskers = 95% CI). **LEFT**: Domain shift evaluation via ten-fold cross-validation for the “TCGA” model (red), “TCGA” model inference on the ZL dataset (green), and ESPWA (purple). ESPWA significantly outperformed the “TCGA” model on the Haitian cohort (accuracy: p = 0.03; AUROC: p = 0.005), highlighting intrinsic technical or biological differences between the two datasets. Regression metrics (PCC, ICC) are provided for ESPWA. **RIGHT**: Validation on an independent paired test set (n = 134) of Haitian breast cancer patients. Parallel slides processed at BWH (blue) and Mirebalais Hospital (orange) demonstrate performance stability across disparate preparation sites and slide scanning hardware, despite marked disparities in WSI quality.

### ESPWA achieves robust performance in ER-status prediction

For binary classification, ESPWA (accuracy: 0.726; 95% CI: 0.723-0.743, AUROC: 0.790; 95% CI: 0.788-0.807) demonstrated improved performance (accuracy: p=0.03; AUROC: p=0.005) compared to the TCGA model in assessing ER status from the ZL cohort (**Figure 2**). ESPWA’s regression model also achieved reasonable performance in predicting quantitative expression of ER, with a PCC of 0.706 (95% CI: 0.660-0.724) and an ICC of 0.524 (95% CI: 0.471-0.551, **Supplemental Figure 1**). To identify subregions of high value within a WSI that ESPWA relied on for classification, we generated exemplar attention heatmaps for both well differentiated and poorly differentiated tumors (**Figure 3A-F**). We observed a high degree of concordance between the high-attention areas highlighted by ESPWA and the tumor regions of interest by an expert pathologist. Furthermore, our decision curve analysis illustrates that ESPWA shows higher net benefit than the current “treat all” paradigm and an alternative “treat none” approach across a wide range of clinician preferences, demonstrating its potential for use in diverse settings (**Figure 3G; Supplemental Figure 2, Supplemental Table 1**).

**Figure 3:**
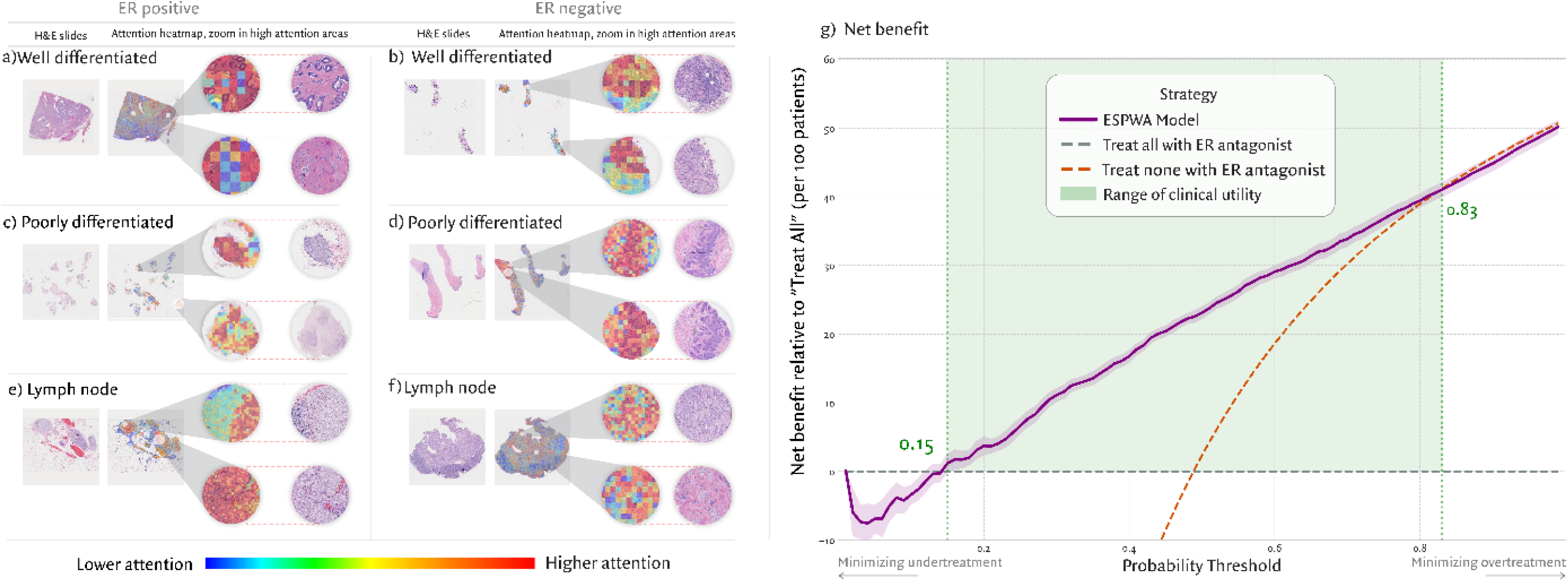
Model introspection. A-F) Exemplar attention heatmaps for H&E WSIs. Red, orange and yellow signify high-attention areas of the H&E image, green, blue and purple signify low-attention areas. G) Decision curve analysis as per Vickers et al^21^ (“interventions avoided” variant) A) This is a well-differentiated ER-positive tumor case (excision specimen), which ESPWA correctly predicted. The upper panel shows gland formation, but this area to a human eye looks like poorly preserved benign glands with nuclear atypia, and part of a fibroadenomatoid background. The lower panel shows distinct nests of infiltrating tumor cells with low-grade nuclei, consistent with a low-grade, well-differentiated tumor (tubule score 3, nuclear score 1, mitoses score 1 in the Bloom-Richardson grading system). B) This is a well-differentiated ER-negative tumor case (biopsy specimen), which ESPWA correctly predicted. In the upper panel, areas of high-attention correlate with tubules (red) in a dense stroma of inflammatory cells (orange and yellow). It is not obvious to the human eye that this is infiltrative tumor. By contrast the lower panel shows a distinct infiltrating tumor with lobular features. There are no tubules, the architecture is single file spread, nuclei are small, and mitoses are absent (tubule score 3, nuclear score 1, mitoses score 1 in the Bloom-Richardson grading system). Normally this low-grade histomorphology would suggest ER-positive status, but in the setting of ER-negative status, this is best classified as a histiocytoid invasive lobular carcinoma. C) This is a poorly differentiated ER-positive tumor case (biopsy specimen), which ESPWA correctly predicted. The upper panel high-attention areas illustrate a solid sheet of tumor cells showing pleomorphic nuclei and frequent mitoses (tubule score 3, nuclear score 3, mitoses score 3 in the Bloom-Richardson grading system). The lower panel of high attention shows necrotic tumor, suggesting high cell turnover. D) This is a poorly differentiated ER-negative tumor case (biopsy specimen), which ESPWA correctly predicted. High-attention areas in the upper and lower panels depict infiltrating solid nests of tumor cells showing pleomorphic nuclei and frequent mitoses (tubule score 3, nuclear score 3, mitoses score 3 in the Bloom-Richardson grading system). E) This is an ER-positive lymph node metastasis (excision specimen), which ESPWA correctly predicted. High-attention areas show tumor cells with uniform, low-grade nuclei and scant mitoses, consistent with low-grade tumor. The lymphocytic background is mostly low attention (green). F) This is an ER-negative lymph node metastasis (excision specimen), which ESPWA correctly predicted. High-attention areas illustrate tumor cells; the upper panel shows pleomorphic nuclei and numerous mitoses, consistent with high-grade tumor. The lower panel highlights pleomorphic tumor cells with necrotic tumor. G) The x-axis shows the model threshold used, derived from the relative preference of avoiding unnecessary treatments for those who will not benefit compared to incorrectly withholding treatment from those who would benefit: on the left, a low threshold is applied to the model output because correctly treating all those who can benefit is considered more important, on the right a high threshold is used because avoiding unnecessary treatment is considered more important. The y-axis shows the net benefit relative to the “treat all” strategy, defined as the number of unnecessary treatments correctly avoided minus the number of treatments incorrectly withheld, weighted by the implied preference ratio. ESPWA show higher net benefit than either the “treat all” or “treat none” strategies across a wide range of preferences, demonstrating its potential for impact in wide-ranging clinical scenarios.

ESPWA’s performance is noteworthy, given the high prevalence of poorly differentiated tumors in the Black population^16^. Accurate ER assessment based on H&E-stained slides of poorly differentiated tumors, due to loss of typical indicators of ER-positive status (e.g., tubule architecture, scant mitoses, low nuclear grade), is a difficult task for pathologists. In the ZL cohort, 65.3% (2254/3448) of tumors were poorly differentiated (**Table 1**), compared to a historical prevalence of 29.7% in the White population. This disparity highlights the increased difficulty of ER classification in the ZL population relative to the TCGA cohort.

**Table 1.** ESPWA Performance Metrics Across Clinical and Histologic Subgroups Across the ZL Cross-Validation Set.

| Subgroup | N. samples | ER+ | ER- | Accuracy | AUROC | AUPRC | PCC | ICC |
| --- | --- | --- | --- | --- | --- | --- | --- | --- |
| Overall |  |  |  |  |  |  |  |  |
| All Samples | 3,448 | 1684 | 1764 | 0.726<br>(0.723-0.743) | 0.790<br>(0.788-0.807) | 0.761<br>(0.752-0.784) | 0.706<br>(0.660-0.724) | 0.524<br>(0.471-0.551) |
| ER Status |  |  |  |  |  |  |  |  |
| ER-positive | 1,684 | 1684 | 0 | 0.723<br>(0.708-0.732) | — | — | — | — |
| ER-negative | 1,764 | 0 | 1764 | 0.729<br>(0.721-0.746) | — | — | — | — |
| ER quantitative expression |  |  |  |  |  |  |  |  |
| ER-Low expression (0-10%) | 144 | 144 | 0 | 0.396<br>(0.303-0.467) | — | — | — | — |
| ER-High expression (>10%) | 1540 | 1540 | 0 | 0.768<br>(0.751-0.776) | — | — | — | — |
| Tumor grade |  |  |  |  |  |  |  |  |
| Grade 1-2 (Well-differentiated) | 1,194 | 749 | 445 | 0.747<br>(0.719-0.765) | 0.775<br>(0.744-0.801) | 0.816<br>(0.785-0.846) | 0.608<br>(0.533-0.625) | 0.423<br>(0.342-0.444) |
| Grade 3 (Poorly - differentiated) | 2,254 | 935 | 1319 | 0.715<br>(0.697-0.737) | 0.776<br>(0.762-0.802) | 0.703<br>(0.675-0.738) | 0.728<br>(0.714-0.761) | 0.546<br>(0.520-0.588) |
| Tissue type |  |  |  |  |  |  |  |  |
| Lymph node | 515 | 265 | 250 | 0.676<br>(0.645-0.694) | 0.707<br>(0.659-0.733) | 0.654<br>(0.625-0.697) | 0.619<br>(0.581-0.730) | 0.437<br>(0.388-0.563) |
| Primary tumor | 2,933 | 1419 | 1514 | 0.735<br>(0.726-0.753) | 0.802<br>(0.793-0.822) | 0.779<br>(0.764-0.803) | 0.721<br>(0.692-0.738) | 0.539<br>(0.504-0.564) |
| Histological subtype |  |  |  |  |  |  |  |  |
| Ductal | 2,102 | 1295 | 807 | 0.710<br>(0.695-0.732) | 0.770<br>(0.750-0.785) | 0.831<br>(0.816-0.844) | 0.714<br>(0.666-0.743) | 0.530<br>(0.474-0.564) |
| Lobular | 138 | 101 | 37 | 0.681<br>(0.639-0.725) | 0.712<br>(0.666-0.769) | 0.860<br>(0.825-0.913) | 0.670<br>(0.540-0.804) | 0.485<br>(0.367-0.648) |
| Other | 1,208 | 288 | 920 | 0.760<br>(0.742-0.773) | 0.806<br>(0.784-0.820) | 0.519<br>(0.484-0.556) | 0.685<br>(0.601-0.708) | 0.511<br>(0.416-0.535) |

### ESPWA is robust to laboratory-specific variability and WSI quality

To evaluate model performance across diverse lab environments, we conducted a paired analysis on a test set of 134 Haitian patients, comparing parallel slides cut, processed, and scanned at BWH (United States) versus Mirebalais Hospital (Haiti). Quantitative assessment via GrandQC revealed a significant increase in technical artifacts in the Mirebalais-prepared WSIs across all four evaluated categories: tissue folds, dark/foreign spots, edge bubbles, and out-of-focus patches (**Figure 4; Supplemental Table 2**).

**Figure 4:**
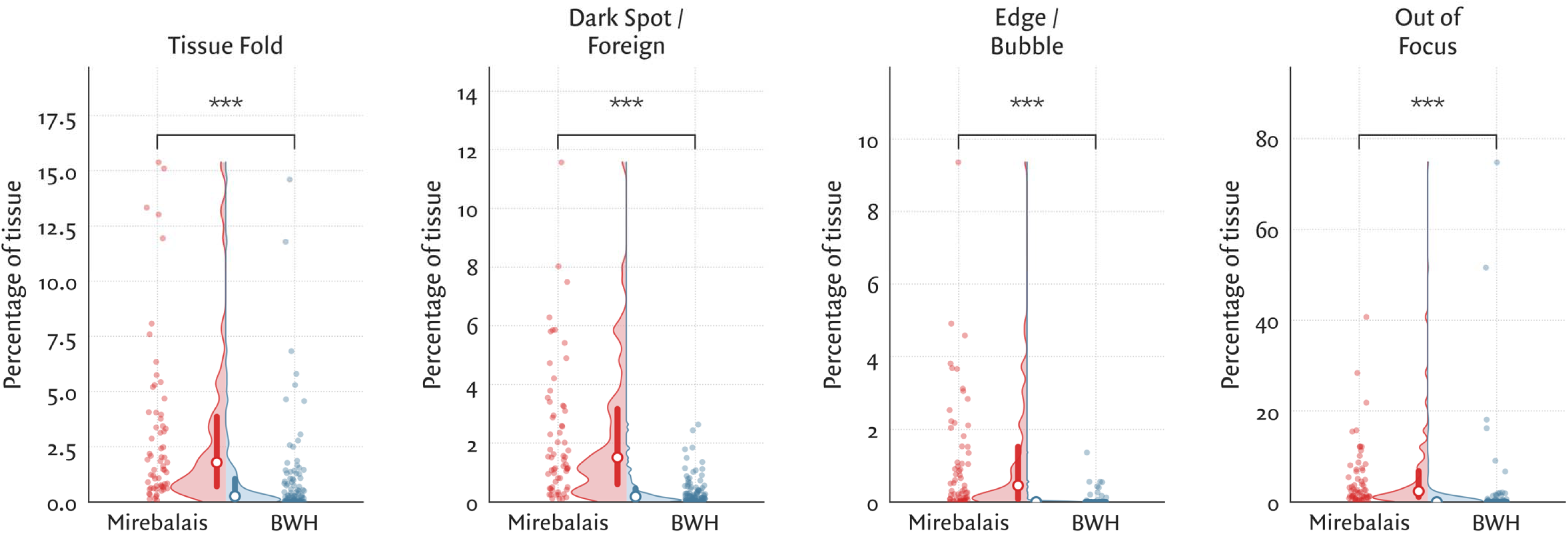
Comparison of WSI artifact burden by preparation site. Scatter and violin plots illustrate the percentage of tissue affected by common histopathological artifacts: (A) Tissue Folds, (B) Dark Spots or Foreign Material, (C) Edge/Bubble, and (D) Out of Focus. Individual data points represent single WSIs, color-coded by site: red for Mirebalais-prepared WSIs and blue for BWH-prepared WSIs. Within each plot, white circles indicate median values and thick vertical bars represent the interquartile range (IQR). Notably, Mirebalais-prepared WSIs exhibited a significantly higher median artifact burden across all evaluated categories compared to BWH-prepared WSIs: tissue folds (1.78% vs 0.26%), dark spots (1.51% vs 0.18%), edge/bubbles (0.46% vs 0.00%), and out of focus regions (2.32% vs 0.02%). Asterisks indicate statistical significance (p < 0.001 for all comparisons).

Despite the significantly higher burden of artifacts in the LMIC-prepared cohort, ESPWA demonstrated notable performance stability. There was no statistically significant drop in performance between the BWH-prepared test set (AUROC: 0.794; 95% CI: 0.696-0.887) and the Mirebalais-prepared test set (AUROC: 0.805; 95% CI: 0.715-0.899; p = 0.753; **Figure 2**). Furthermore, these results were consistent with the ten-fold cross-validation performance with the broader ZL cohort (AUROC: 0.790; 95% CI: 0.788-0.807), with no significant differences when compared to either the BWH-prepared (p = 0.204) or Mirebalais-prepared (p = 0.415) test sets. This parity in performance across institutions, despite marked differences in slide quality and scanning hardware, suggests that ESPWA is invariant to the technical variability and quality issues inherent in Haitian and LMIC pathology workflows.

### Subgroup analyses

Next, we conducted an analysis across clinically relevant subgroups within our cohort (**Table 1**). There was no difference in ESPWA’s ability to assess ER status between the ER-positive (0.723, 95% CI: 0.708-0.732) and ER-negative subgroups (0.729, 95% CI: 0.721-0.746, p=0.705). Within the ER-positive cohort, there was a marked difference in ESPWA accuracy between the ER-low positive (1-10% by IHC; 0.396, 95% CI: 0.303-0.467), and ER-high positive cohorts (>10%; 0.768, 95% CI: 0.751-0.776, p<0.001). This discrepancy accurately reflects tumor biology, as ER-low positive tumors behave more similarly to ER-negative tumors, which are resistant to ET.

Next, we compared ESPWA’s performance between well-differentiated (grade 1-2) and poorly differentiated (high-grade) tumors. While the AUROCs were comparable between these two groups, the AUPRC was higher for grade 1-2 tumors (0.816, 95% CI: 0.785-0.846) than for grade 3 tumors (0.703, 95% CI: 0.675-0.738, **Supplemental Figure 3**). This reflects the higher prevalence of ER-positive tumors in the grade 1-2 cohort (749/1194 = 62.7%) compared to the grade 3 cohort (935/2254 = 41.4%), leading to a higher baseline precision value. ESPWA’s observed AUPRC for grade 3 tumors reflects reasonable performance above random chance in a subgroup that is challenging for human experts.

Next, ESPWA’s performance was higher on primary tumor samples (AUROC = 0.802; 95% CI: 0.793-0.822), compared to lymph node metastases (AUROC = 0.707, 95% CI: 0.659-0.733, p=0.005). This phenomenon may be due to the presence of dense lymphocytic infiltrates in lymph nodes, which are associated with ER-negative status in primary tumors but not in lymph node metastases. Finally, we did not observe substantial variation in performance across different histological subtypes. Confusion matrices for all subgroup analyses (**Supplemental Figure 4**) show no major imbalance between false positives and false negatives across any subgroup.

### ESPWA outperforms a human expert in ER-status prediction

To benchmark model performance against human expertise, we compared ESPWA’s classification accuracy to that of an academic breast pathologist with 22 years of experience (J.E.B.). The human expert evaluated ER status based on tumor morphology on H&E WSIs from the ZL cohort and obtained an accuracy of 0.639 (95% CI: 0.619-0.652). By comparison, ESPWA achieved a superior accuracy of 0.726 (95% CI: 0.723-0.743, p<0.001) across the entire cohort (**Figure 5, Supplemental Table 3**) and across most subgroups.

**Figure 5:**
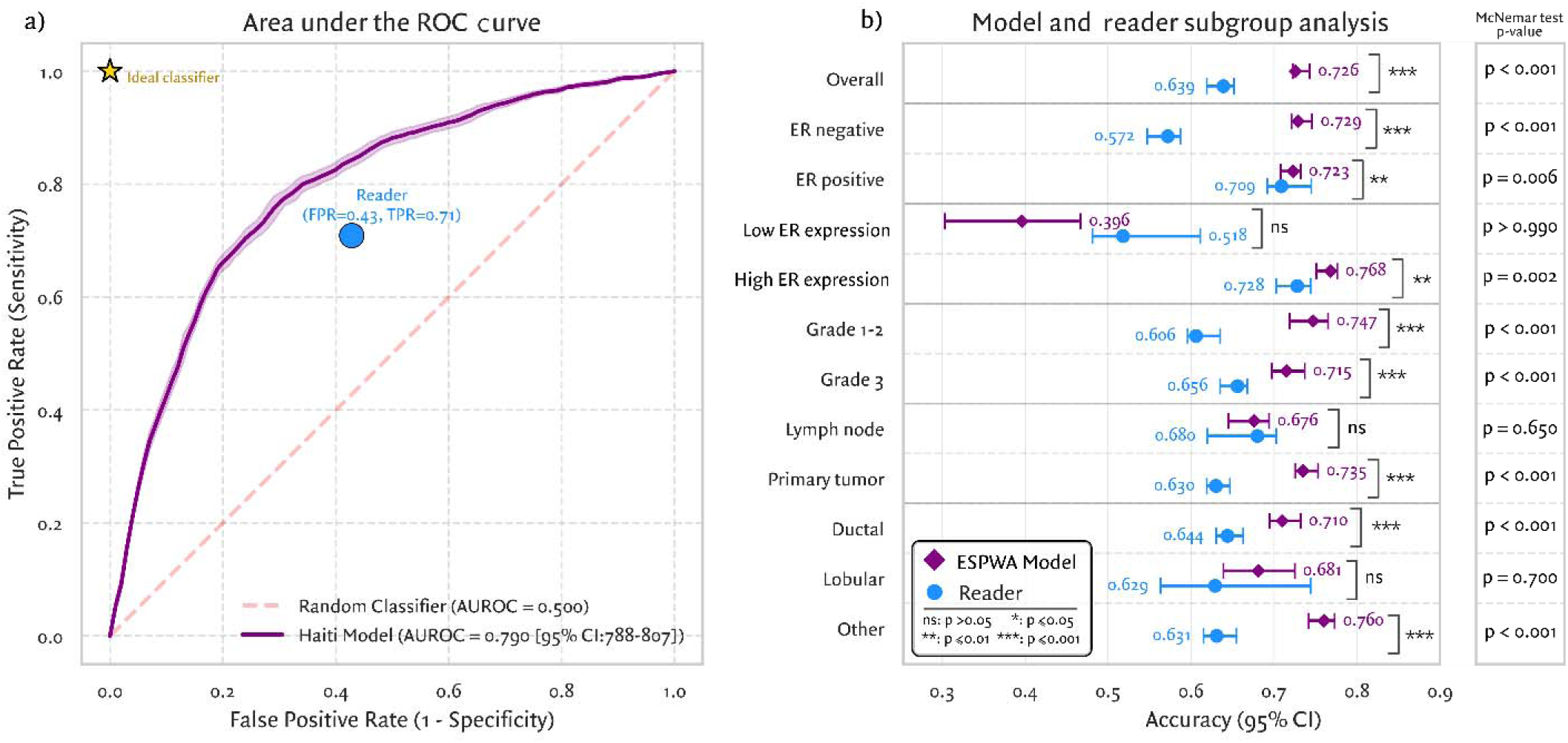
Comparison of ESPWA with a breast pathologist in ER status prediction from H&E WSI. A) Receiver operating characteristic (ROC) curve depicting performance of ESPWA (purple line) compared to a breast pathologist (Reader, blue dot). The ideal classifier is represented by the yellow star at the top left corner. B) Subgroup analysis comparing classification accuracy between ESPWA (purple) and the breast pathologist (blue circles) across clinical and pathological subgroups. Error bars represent 95% confidence intervals. Statistical comparisons by McNemar tests for all subgroups are provided.

## DISCUSSION

DL, through its ability to execute tasks involving high-dimensional data, is poised to transform healthcare delivery in resource-limited settings. In recent years, DL has enabled the development of tools that analyze clinical data to provide a timely diagnosis across diverse diseases^24–26^ in LMICs. However, to date, few of these tools have extended beyond diagnostic applications to inform *therapeutic* decision making. While diagnostic tools are an essential part of clinical care, tools that interrogate an individual patient’s disease biology to guide selection of the *optimal* treatment are equally critical to improving cancer outcomes.

Here, we present ESPWA, a DL model that uses widely available H&E WSIs to guide precision-based use of ET in LMIC breast cancer patients. By recognizing spatial patterns of therapeutically relevant cell populations within the tumor microenvironment, ESPWA achieved strong performance in ER status prediction across clinical and histologic subgroups. Our model performed well in poorly differentiated tumors – a clinically noteworthy finding given the high prevalence of such tumors in LMICs with predominantly Black populations^9^. Moreover, ESPWA outperformed an academic breast pathologist in assessing ER status directly from H&E WSIs, suggesting that DL may uncover biologically meaningful patterns on H&E WSIs that are not readily discernable by pathologists. Therefore, in settings where IHC is inaccessible and empiric ET is the standard of care, ESPWA offers a precision-based approach that could significantly reduce the administration of ineffective and toxic therapies.

A defining strength of this study is the demonstration of ESPWA’s generalizability across disparate laboratory environments. DL models in the medical domain are notoriously sensitive to “domain shift”, as performance often degrades when encountering the “noise” of different staining protocols or scanning hardware. We addressed this potential limitation through a paired analysis of 134 Haitian patients, whose tissue was processed in parallel at BWH (HIC setting) and Mirebalais Hospital (LMIC setting). Despite a significantly higher burden of technical artifacts in the Mirebalais-prepared WSIs (e.g., tissue folds, out-of-focus WSI patches), ESPWA maintained remarkable performance stability. The lack of a statistically significant drop in AUROC between these two cohorts and the internal cross-validation cohort suggests that ESPWA is invariant to the technical heterogeneity and quality disparities inherent in LMIC pathology workflows. Furthermore, our use of label-preserving augmentation during training likely enhanced ESPWA’s ability to generalize across different lab contexts. Such robustness is a prerequisite for real-world deployment to ensure that clinical insights remain reliable across a range of WSI quality conditions.

Furthermore, our results underscore the necessity of domain-specific training. We illustrated that models trained exclusively on HIC datasets (e.g., TCGA) fail to generalize to the Haitian population. This phenomenon likely stems from a combination of distinct breast cancer biology^16,17^ and differences in tissue processing and fixation quality between HICs and LMICs^14,27^. By using a large training set of over 3400 H&E WSIs from Haitian patients, we ensured that ESPWA was adapted to the biological and technical nuances of the population it is intended to serve. Consequently, ESPWA and our ZL dataset provide a valuable foundation for future efforts aimed at building deployable precision oncology tools for the LMIC setting.

Given the limited infrastructure in LMICs, any deployed tool must be cost-effective and technologically appropriate. The American Society for Clinical Pathology (ASCP) has led diagnostic capacity-building initiatives across multiple LMICs across Africa, Asia, and the Caribbean, supporting the development of foundational capabilities such as slide digitization and secure file transfer. Within such environments, ESPWA could be deployed using a cloud server with minimal additional resources.

Based on this scalable framework, prospective feasibility studies have been initiated across ASCP partner sites in sub-Saharan Africa to evaluate the operational integration and clinical utility of ESPWA. These efforts will generate additional data to refine ESPWA, assess its performance in real-world settings, and extend model capabilities to derive progesterone receptor (PR) and HER2 status from H&E WSIs. These molecular markers are therapeutically relevant, as ER-positive, HER2-positive patients typically respond less favorably to ET compared to ER-positive, HER2-negative patients. Incorporating PR status may also enhance model performance through regularization, given the frequent co-expression of ER and PR.

We acknowledge that ESPWA demonstrated reduced performance in certain subgroups, particularly the ER-low positive cohort. This observation was expected, as these tumors often behave biologically like ER-negative tumors ^28,29^. However, since the primary objective of ESPWA was to identify patients most likely to benefit from ET, misclassifying ER-low positive tumors as ER-negative tumors may actually align with its intended therapeutic utility. Additionally, we observed that ER classification of lymph node metastases was less accurate compared to that of the overall dataset. Notably, the inclusion of lymph node metastases in our training set is a strength, as most prior DL models in breast cancer have been trained exclusively on WSIs of primary tumors. Given the distinct histological differences between lymph node metastases (e.g., dense lymphocyte-rich background) and primary tumors (e.g., epithelial or glandular tissue infiltrating fibrous or fatty stroma), the morphological features predictive of ER status likely vary between these tissue types. Future iterations of ESPWA incorporating larger training sets of lymph node tissue will likely improve performance for this subgroup.

In summary, we present ESPWA, a DL model designed to enable precision-based use of ETs in resource-limited settings. By achieving robust performance across different laboratory environments and varying WSI quality, ESPWA demonstrates that individualized therapeutic insights can be extracted directly from routine H&E WSIs. This scalable, technologically appropriate solution leverages existing infrastructure in LMICs to markedly improves upon the current standard of empiric ET, thus bridging a critical gap in the diagnostic-therapeutic pathway. Our findings add to the growing body of work^6,14,30,31^ in illustrating the promise of DL to democratize precision oncology. Based on this work, we are launching prospective trials to deploy ESPWA across LMICs. Ultimately, we believe this work serves as a blueprint to employ DL to uncover actionable biology within routine WSIs and deliver precision-based care to vulnerable patient populations in low-resource settings.

---

**Supplemental Figure 1:**
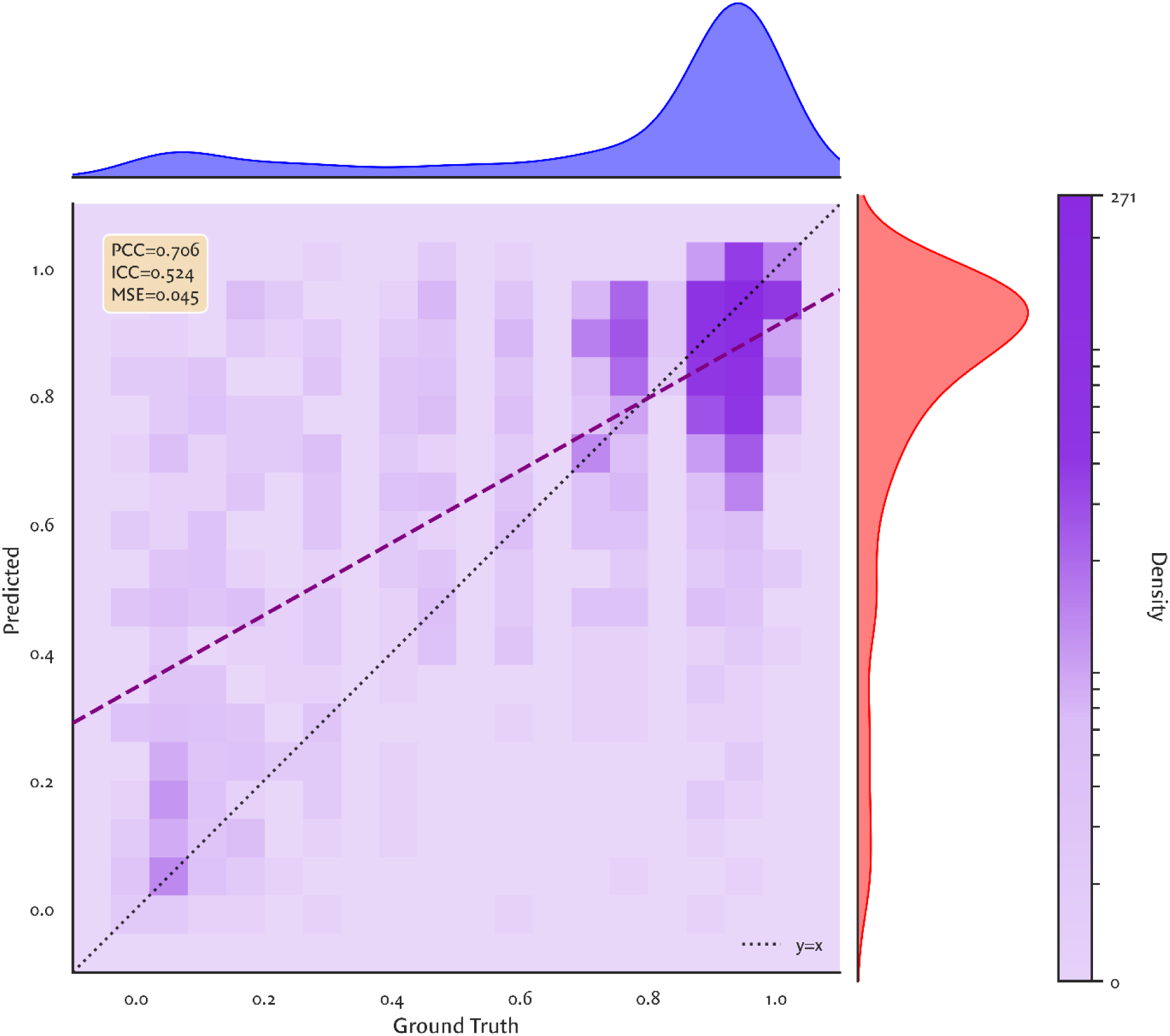
Density plot of predicted vs. ground truth quantitative ER expression values for ESPWA. The heatmap intensity (purple) reflects number of WSI (sample density), with marginal density distributions shown for the ground truth (top, blue) and predicted values (right, red). The purple dashed line indicates the linear regression fit, while the black dotted line represents the ideal y = x relationship. Performance metrics including PCC (0.706), ICC (0.524), and mean squared error (MSE = 0.045) are displayed, illustrating moderat agreement between ESPWA’s predicted and actual ground truth ER expression levels.

**Supplemental Figure 2:**
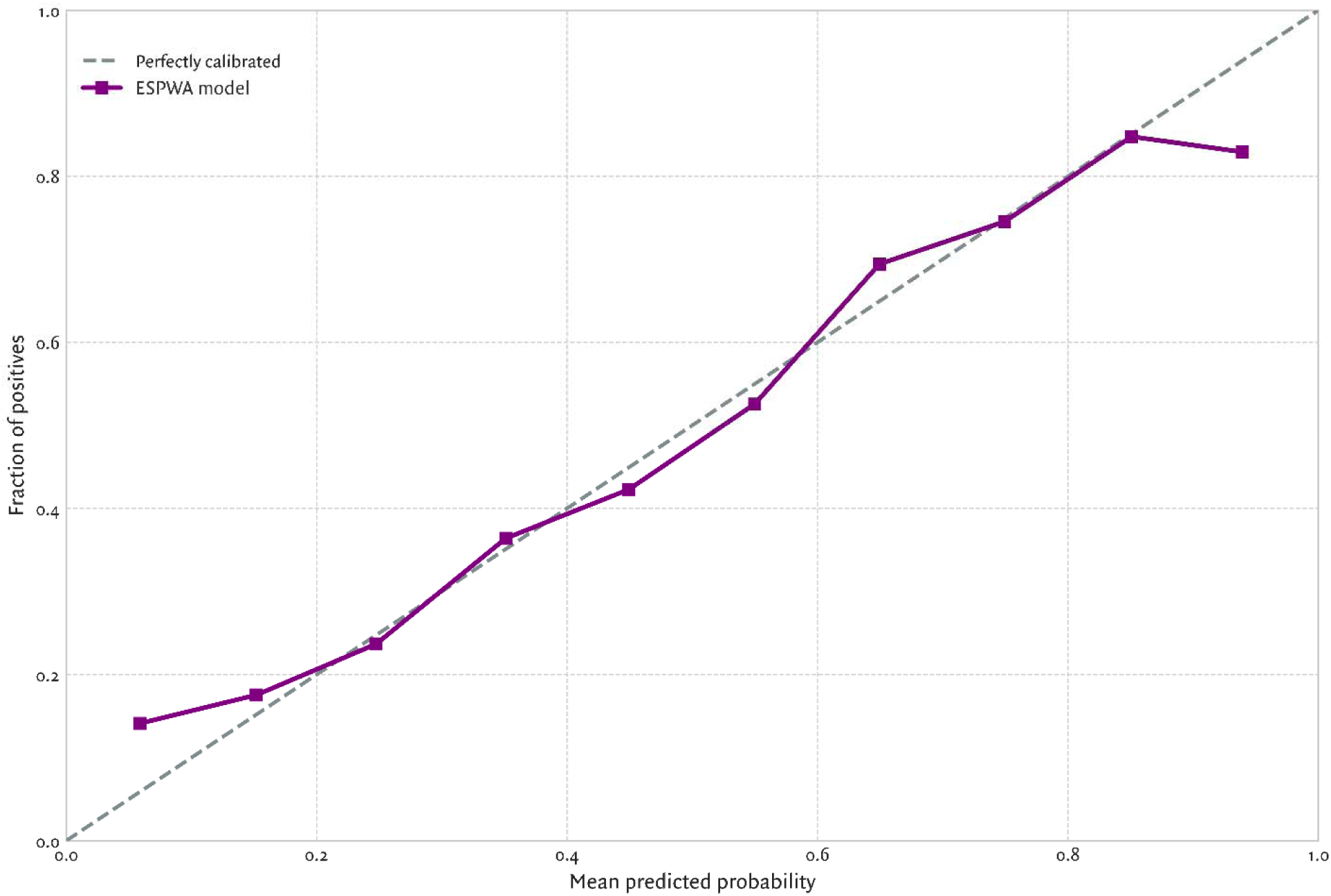
Calibration plot for ESPWA for ER status prediction. Calibration performance of ESPWA (purple) is illustrated by plotting observed fraction of ER-positive cases against the mean predicted probability (model output). This plot illustrates that ESPWA is a well-calibrated model, as this model is in close alignment with a perfectly calibrated model (dashed line), thus enabling our decision curve analysis.

**Supplemental Figure 3:**
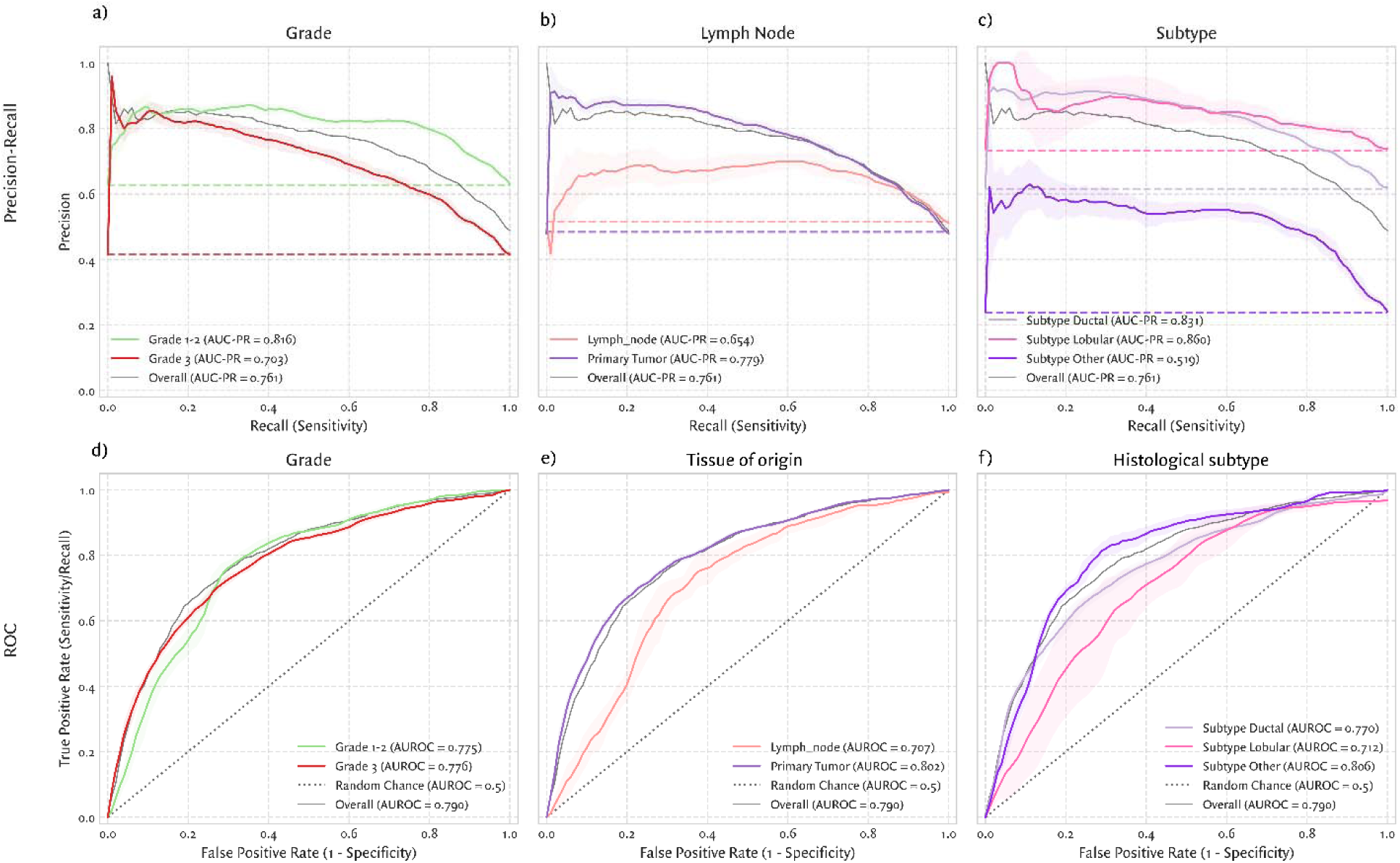
Precision-recall (top row) and ROC curves (bottom row) evaluating model performance stratified by clinical and histologic subgroups. A & B) Precision-recall and AUROC curves for the tumor grade subgroups. C & D) Precision-recall and AUROC curves for the tissue of origin subgroups. E & F) Precision-recall and AUROC curves for histological subtype subgroups

**Supplemental Figure 4:**
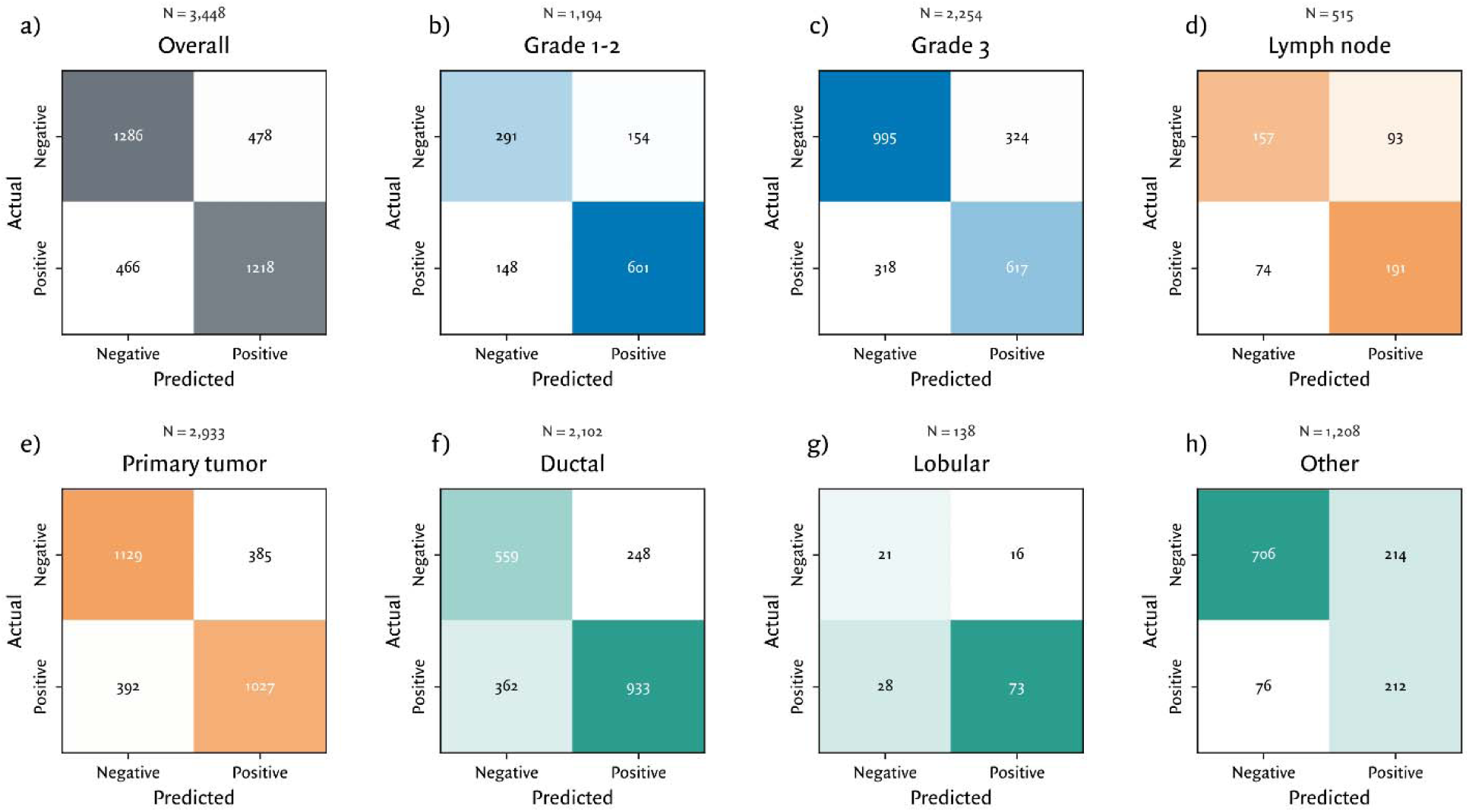
Confusion matrices for ESPWA stratified by clinical and histologic subgroups. (A) overall, (B-C) grade, (D-E) tissue type, (F-H) histologic subtype.

**Supplemental Table 1.** Threshold-based performance and net benefit analysis of ESPWA.

| Cutpoint Strategy | Cutpoint | Sensitivity | Specificity | PPV | NPV | ER positives untreated (n = 1,684) | ER negatives untreated (n = 1,764) | Net benefit relative to “Treat All” (per 100 patients) |
| --- | --- | --- | --- | --- | --- | --- | --- | --- |
| Sensitivity 10% | 0.899 | 10.5%<br>(9.3% - 11.8%) | 98.0%<br>(97.6% - 98.4%) | 83.5%<br>(79.6% - 87.3%) | 53.4%<br>(51.6% - 54.9%) | 1,507<br>(1484 - 1525) | 1,729<br>(1698 - 1767) | 45.3<br>(44.0 - 46.3) |
| Sensitivity 20% | 0.845 | 20.2%<br>(19.1% - 21.2%) | 96.7%<br>(96.2% - 97.8%) | 85.4%<br>(82.7% - 89.6%) | 55.9%<br>(54.6% - 56.8%) | 1,344<br>(1297 - 1403) | 1,706<br>(1646 - 1758) | 42.3<br>(40.1 - 43.7) |
| Default | 0.5 | 72.3%<br>(69.9% - 74.5%) | 72.9%<br>(71.3% - 74.9%) | 71.8%<br>(70.7% - 73.2%) | 73.4%<br>(71.5% - 74.6%) | 466<br>(432 - 470) | 1,286<br>(1253 - 1329) | 23.8<br>(22.2 - 25.1) |
| Optimal (Youden’s J) | 0.482 | 74.5%<br>(72.9% - 74.6%) | 71.5%<br>(70.4% - 72.7%) | 71.4%<br>(70.1% - 72.8%) | 74.6%<br>(73.1% - 74.7%) | 430<br>(399 - 456) | 1,262<br>(1227 - 1306) | 23.2<br>(22.1 - 24.7) |
| Sensitivity 80% | 0.415 | 80.0%<br>(78.7% - 81.2%) | 63.7%<br>(62.6% - 65.6%) | 67.8%<br>(66.6% - 70.0%) | 77.0%<br>(75.9% - 78.5%) | 336<br>(305 - 362) | 1,123<br>(1076 - 1155) | 18.8<br>(16.9 - 21.0) |
| Sensitivity 90% | 0.252 | 90.0%<br>(89.2% - 91.1%) | 43.1%<br>(41.4% - 44.8%) | 60.2%<br>(59.2% - 62.2%) | 81.9%<br>(80.0% - 83.7%) | 168<br>(156 - 185) | 761<br>(743-786) | 7.6<br>(4.7 - 9.5) |
| Treat All (baseline) | 0.000 | 100%<br>(100% - 100%) | 0%<br>(0% - 0%) | 48.8%<br>(47.2% - 50.4%) | 0%<br>(0% - 0%) | 0<br>(0 - 0) | 0<br>(0 - 0) | 0<br>(0 - 0) |

This table presents ESPWA’s performance on the validation cohort under various model thresholds. For each metric, we report the point estimate and 95% confidence interval (CI), calculated via a bootstrap procedure with 1,000 resamples.
The column “Cutpoint Strategy” describes the criterion for selecting the classification threshold, which may include the point of maximum accuracy (Youden's Index), thresholds that ensure predefined sensitivity levels (e.g., 10%, 20%, 80%, 90%), and a comparison to the “Treat All” standard of care, corresponding to the highest achievable sensitivity. The “Cutpoint” column indicates the numerical value of the model's probability threshold, with scores greater than or equal to this value classified as ER-positive.
“Sensitivity” measures the proportion of ER-positive cases correctly identified, calculated as True Positives divided by the sum of True Positives and False Negatives, while “Specificity” measures the proportion of ER-negative cases correctly identified, calculated as True Negatives divided by the sum of True Negatives and False Positives.
“Positive Predictive Value (PPV)” indicates the probability that a case classified as ER-positive is truly positive, calculated as True Positives divided by the sum of True Positives and False Positives, and “Negative Predictive Value (NPV)” indicates the probability that a case classified as ER-negative is truly negative, calculated as True Negatives divided by the sum of True Negatives and False Negatives.
The column “ER positives untreated” represents the number of ER-positive cases incorrectly classified as negative (False Negatives), corresponding to patients who would be missed for treatment; the cohort includes 1,684 ER-positive patients.
The column “ER negatives untreated” represents the number of ER-negative cases correctly classified as negative (True Negatives), corresponding to patients who would correctly avoid unnecessary treatment; the cohort includes 1,764 ER-negative patients. Finally, “Net Benefit” measures the clinical utility of the model by weighing the benefits of true positives against the harms of false positives, with higher values indicating greater usefulness compared to the default strategies of treating all or no patients.

**Supplemental Table 2.** Technical artifact burden in the independent test set (n = 134).

| Artifact Category<br>(% within WSI) | Preparation Site | Median (Interquartile<br>Range) | Mean (Standard<br>Deviation) |
| --- | --- | --- | --- |
| Tissue Folds | Mirebalais Hospital | 1.78% (0.70-3.86%) | 3.09% (3.66%) |
|  | Brigham &<br>Women's Hospital | 0.26% (0.03-1.05%) | 1.04% (2.20%) |
| Dark Spots or<br>Foreign Material | Mirebalais Hospital | 1.51% (0.60-3.18%) | 2.26% (2.27%) |
|  | Brigham &<br>Women's Hospital | 0.18% (0.05-0.46%) | 0.36% (0.50%) |
| Edges or Bubbles | Mirebalais Hospital | 0.46% (0.08-1.52%) | 1.13% (1.62%) |
|  | Brigham &<br>Women's Hospital | 0.00% (0.00-0.00%) | 0.05% (0.18%) |
| Out of Focus | Mirebalais Hospital | 2.32% (1.05-6.82%) | 5.01% (7.09%) |
|  | Brigham &<br>Women's Hospital | 0.02% (0.00-0.22%) | 2.05% (9.46%) |

**Supplemental Table 3.** Pathologist Performance Metrics Across Clinical and Histologic Subgroups Across the ZL dataset.

|  | Subgroup | N. samples | ER+ | ER- | Accuracy | Precision | Recall | Sensitivity | Specificity |
| --- | --- | --- | --- | --- | --- | --- | --- | --- | --- |
| Overall |  |  |  |  |  |  |  |  |  |
|  | All Samples | 3,448 | 1684 | 1764 | 0.639<br>(0.619-0.652) | 0.611<br>(0.595-0.622) | 0.709<br>(0.677-0.736) | 0.709<br>(0.677-0.736) | 0.572<br>(0.562-0.589) |
| ER Status |  |  |  |  |  |  |  |  |  |
|  | ER-positive | 1,684 | 1684 | 0 | 0.709<br>(0.692-0.745) | — | — | — | — |
|  | ER-negative | 1,764 | 0 | 1764 | 0.572<br>(0.547-0.587) | — | — | — | — |
| ER quantitative expression |  |  |  |  |  |  |  |  |  |
|  | ER-Low<br>expression (0-10%) | 144 | 144 | 0 | 0.518<br>(0.481-0.611) | — | — | — | — |
|  | ER-High<br>expression (>10%) | 1,479 | 1475 | 0 | 0.728<br>(0.703-0.744) | 0.999<br>(0.996-1.000) | 0.728<br>(0.703-0.745) | 0.728<br>(0.703-0.745) | 0.750<br>(0.113-1.000) |
| Tumor grade |  |  |  |  |  |  |  |  |  |
|  | Grade 1-2<br>(Well-differentiated) | 1,194 | 749 | 445 | 0.606<br>(0.596-0.635) | 0.655<br>(0.630-0.677) | 0.842<br>(0.826-0.878) | 0.842<br>(0.826-0.878) | 0.158<br>(0.143-0.189) |
|  | Grade 3 (Poorly<br>- differentiated) | 2,254 | 935 | 1319 | 0.656<br>(0.635-0.668) | 0.567<br>(0.535-0.617) | 0.597<br>(0.583-0.605) | 0.597<br>(0.583-0.605) | 0.695<br>(0.661-0.728) |
| Tissue type |  |  |  |  |  |  |  |  |  |
|  | Lymph node | 515 | 265 | 250 | 0.680<br>(0.620-0.703) | 0.696<br>(0.656-0.728) | 0.693<br>(0.632-0.731) | 0.693<br>(0.632-0.731) | 0.665<br>(0.599-0.699) |
|  | Primary tumor | 2,933 | 1419 | 1514 | 0.630<br>(0.619-0.647) | 0.595<br>(0.583-0.612) | 0.713<br>(0.693-0.738) | 0.713<br>(0.693-0.738) | 0.554<br>(0.533-0.577) |
| Histological subtype |  |  |  |  |  |  |  |  |  |
|  | Ductal | 2,102 | 1295 | 807 | 0.644<br>(0.630-0.663) | 0.709<br>(0.692-0.737) | 0.689<br>(0.669-0.699) | 0.689<br>(0.669-0.699) | 0.576<br>(0.553-0.633) |
|  | Lobular | 138 | 101 | 37 | 0.629<br>(0.563-0.744) | 0.701<br>(0.631-0.840) | 0.844<br>(0.766-0.914) | 0.844<br>(0.766-0.914) | 0.080<br>(0.034-0.184) |
|  | Other | 1,208 | 288 | 920 | 0.631<br>(0.615-0.655) | 0.401<br>(0.358-0.443) | 0.748<br>(0.717-0.777) | 0.748<br>(0.717-0.777) | 0.588<br>(0.563-0.622) |

## Acknowledgments

We thank Scott Kilcoyne of DigitCells for the use of a KFBIO KF-PRO-005 scanner, and appreciate helpful feedback from Gromit Persimmon Perrino.

## Data Sharing Statement

Any requests for raw and analyzed data will be reviewed by the Massachusetts General Hospital Institutional Review Board (IRB). Any data and materials (e.g. imaging or IHC data) that can be shared will need approval from the MGH IRB and a Material Transfer Agreement in place. All data shared will be deidentified.

## Code Availability

Code from this study is available at: https://github.com/QTIM-Lab/ESPWA.

## Acknowledgments of Research Support

This work was supported by the intramural research program (ZIACP101237-01) of the US National Cancer Institute/NIH/DHHS, the Department of Defense Breast Cancer Breakthrough Award – Level 2 (BC241017), William G. Kaelin, Jr., M.D., Physician-Scientist Award of the Damon Runyon Cancer Research Foundation (PST-36-21), American Association for Cancer Research Breast Cancer Research Fellowship (21-40-49-KIM), American Brain Tumor Association Basic Research Fellowship In Honor of Paul Fabbri, American Society of Clinical Oncology/Conquer Cancer Young Investigator Award, American Academy of Neurology Career Development Award, American Cancer Society Institutional Research Grant (IRG-21-130-10), NCI K12CA090354, and NIBIB K08EB037077.

