## Supplemental Methods for "ESPWA: a deep learning tool to inform precision-based use of endocrine therapy in resource-limited settings"

**SUPPLEMENTARY METHODS**

**Whole-Slide Image Processing**

For both the TCGA and ZL datasets, H&E-stained slides were digitized at 40x magnification using a KFBIO KF-PRO-005 scanner. We performed quality control and tissue segmentation using the HEST library's segmentation model [1], a fine-tuned DeepLabV3 architecture with a ResNet50 backbone. This process generated a tissue mask to exclude background, scanning artifacts, and pen markings. Using this mask, each WSI was partitioned into non-overlapping 256x256 pixel patches at 20x magnification.

To augment the training data, we implemented two label-preserving strategies designed to enhance model generalization by simulating common sources of variability in histopathology.

1. **HED Color Augmentation**: To address the significant color and intensity variations that arise from differences in tissue preparation and staining between batches, we applied procedural color perturbations at the tile level. This technique, a standard approach for improving model robustness to domain shift, involves first converting RGB patches into the Hematoxylin-Eosin-DAB (HED) color space. This decouples the image into its constituent stain components, allowing for targeted modifications. We then adjusted the intensity of the hematoxylin and eosin channels by a factor sampled uniformly from a range of -0.2 to 0.2. This range is consistent with established methods for simulating realistic stain variations in histopathology, chosen to generate a spectrum of diverse appearances without introducing significant artifacts that could negatively impact learning [3]. This process was used to pre-compute four distinct augmented versions of each slide.
2. **Tile Shifting Augmentation**: To improve the model's spatial invariance and robustness to minor variations in the digitization process, we implemented a tile-shifting strategy. This method modifies the starting coordinates of the tiling grid applied to each WSI. We generated four additional sets of tiles by shifting the grid by half a patch's length (128 pixels) horizontally, vertically, and in both diagonal directions. This creates new patch configurations with slightly different spatial contexts, encouraging the model to learn more generalizable morphological features.

During the training loop, we employed a two-stage probabilistic sampling scheme to introduce feature variability on-the-fly, balancing the need for data diversity with the computational constraints of MIL workflows that rely on pre-computed features. First, at the slide level, each slide had a 20% probability of being selected for augmentation. If a slide was chosen, a second sampling stage was initiated at the tile level, where each individual tile within that slide had a 50% chance of having its feature vector replaced. For each tile selected for replacement, one of its corresponding pre-computed augmented feature vectors was chosen uniformly at random. This stochastic process ensures that the model is exposed to a diverse and continually changing combination of augmented features during training.

**Feature Extraction**

From each 256x256 pixel tissue patch, we extracted a 768-dimensional feature vector using a frozen, pre-trained CONCH v1.5 image encoder model. These features were stored in HDF5 format for efficient retrieval during model training.

**Model Architecture**

We employed a single-branch CLAM architecture (CLAM-SB) with a “small” size configuration [2]. The model's first step involves a feature transformation: the input bag of 768-dimensional patch features from the CONCH encoder is passed through a fully connected layer with a ReLU activation function. This layer projects the features into a 512-dimensional latent space. This intermediate representation is then processed by a gated attention network, which computes attention scores for each patch. These scores are used to produce a weighted-average, 512-dimensional slide-level feature vector, which is finally passed to a prediction head.

For the quantitative prediction of ER percentage, we developed a regression variant of the architecture. In the standard classification model, the prediction head is a fully-connected linear layer that performs a linear mapping from the 512-dimensional slide-level feature vector to a 2-dimensional output space, representing the logits for the binary classification task (ER-positive vs. ER-negative).

For the regression task, we replaced this classification layer with a new, single-output regression head. This head consists of a single linear neuron that performs a linear transformation from the 512-dimensional slide-level feature vector to a single scalar value. No final activation function (e.g., sigmoid) is applied to this output, resulting in an unbounded prediction appropriate for a regression task.

**Model Training**

For both models, we performed 10-fold cross-validation with splits stratified by patient to prevent data leakage. Each fold was trained for a maximum of 200 epochs using the Adam optimizer (initial learning rate 1 × 10⁻⁴, weight decay 1 × 10⁻⁵) and a batch size of one WSI.

The loss function for the classification model combined a slide-level (bag) cross-entropy loss with an instance-level loss designed to refine the attention mechanism. For the instance-level component, we employed a top-k sampling strategy based on attention scores [2]. For each slide, this sampler selected the k=8 patches with the highest attention scores (treated as positive instances) and the k=8 patches with the lowest attention scores (treated as negative instances). This subset of 2k patches was used to compute the instance-level cross-entropy loss. The total loss was a weighted combination of these two components (total_loss = 0.7 * bag_loss + 0.3 * instance_loss), following established methods that have demonstrated robust performance [2]. To address class imbalance at the slide level, we employed a class-weighted sampling strategy.

For the regression model, the ground truth ER percentage scores were normalized to a [0, 1] range prior to training for compatibility with the MSE loss function. As a direct consequence of this modification, the instance-level clustering supervision mechanism, a key component of the CLAM classification framework, was removed as it is conceptually incompatible with a slide-level continuous target. The resulting regression model is therefore optimized end-to-end using only the slide-level MSE loss.

To prevent overfitting, we incorporated a dropout rate of 0.25 and an early stopping criterion with a patience of 20 epochs, monitoring the validation loss. Early stopping was not initiated before a minimum of 25 training epochs had been completed.

**Computational Environment and Reproducibility**

Model training was conducted on a SLURM-managed high-performance computing (HPC) cluster, using single NVIDIA RTX 8000 GPUs with 4 CPU cores and 200 GB of RAM. The data loading pipeline utilized 4 parallel workers to ensure efficient data throughput. During training, data order was shuffled at each epoch to improve generalization. For validation and testing, data was processed in a fixed, sequential order to ensure deterministic evaluation.

To ensure reproducibility, we used a fixed random seed and enabled deterministic algorithm behavior in the software libraries.
